# Fixation time in evolutionary graphs: a mean field approach

**DOI:** 10.1101/391508

**Authors:** Mahdi Hajihashemi, Keivan Aghababaei Samani

## Abstract

A novel analytical method is proposed for calculation of average fixation and extinction times of mutants in a general structured population of two types of species. The method is based on Markov chains and uses a mean field approximation to calculate the corresponding transition matrix. Analytical results are compared with the results of simulation of the Moran process on a number of population structures.

---

Evolutionary Graph Theory (EGT) [1] is one of the most celebrated methods to study the evolution of species in structured populations. In this theory one considers a constant-size population of individuals which are connected to each other through a (directed) network which is called evolutionary graph [2]. A *fitness* is assigned to each type of species. At each time step one individual is selected for reproduction with a probability proportional to its fitness. Then it puts its offspring in place of one of its neighbors with a probability determined by evolutionary graph. This is in fact a generalization of the so called *Moran Process* [3] which takes place in a structured population instead of a well-mixed population.

This theory has been vastly studied in recent years and different features and generalizations of it are addressed. Here we confine ourselves to populations constructed from two types of species whom we call *residents* and *mutants*. An interesting process is to start with just one mutant and see what the fate of the system is. In fact, the system will end up in one of the two possible states, namely, *fixation* or *extinction* of mutants. Two main quantities corresponding to this process are *fixation probability* and *fixation time*. Fixation probability is the probability for a single mutant to take over the whole population and fixation time is the average (conditional) time needed for this result. Both of these quantities are investigated by many researchers [4–9]. Obtaining fixation time is more challenging than fixation probability. In this letter we introduce a novel analytical method to find conditional fixation and extinction times and use it to obtain fixation time on a random graph in mean field approximation.

## Method

Consider a graph with *N* nodes. Each node can be one of two types, namely residents and mutants. Each type has its own fitness which is 1 for residents and r for mutants. A Moran process is running on top of this graph. At each time step one node is selected for reproduction with a probability proportional to its fitness. Then one of its neighbors is selected randomly and is replaced by the reproduced offspring. There are two important quantities corresponding to this process. The first one is fixation probability, i. e. the probability for a single mutant to take over the whole network. The second one is the average time for a single mutant to take over the whole network.

Here we introduce a mean-field approach to calculate these two quantities for a category of complex networks using Markov chains.

There is a Markov chain corresponding to the above process. This Markov chain is shown in Fig. 1. Each *state* of this chain is specified by the number of mutants. The state *i* represents all configurations of evolutionary graph with *i* mutants. The transition matrix of this Markov chain is defined as follows:

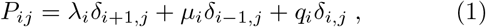

or in matrix form

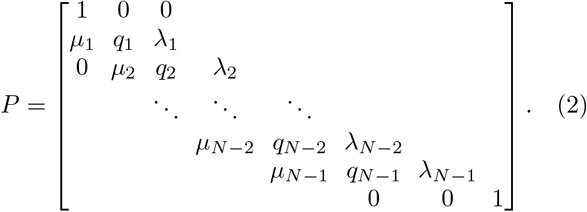

Clearly, λ*_i_* + μ*_i_* + q*_i_* − 1 and λ_0_ — *μ*_0_ − λ_N_ − *μ_N_* − 0. States *S*_0_ and *S_N_* are absorbing states. The first one corresponds to extinction of mutants and the second one corresponds to their fixation. In fact, λ*_i_* is the (average) probability of increasing the number of mutants from *i* to *i* + 1 and *μ_i_* is the (average) probability of decreasing this number from *i* to *i* − 1. To calculate λ*_i_* and *μ_i_* exactly, one has to take into account all graph configurations with i, i+1, and *i* − 1 mutants. Generally, this is not an easy task, but as we will show one can obtain them approximately for a number of graph topologies.

**FIG. 1.**
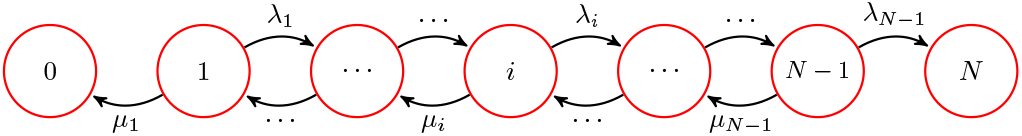
(color on line) Markov chain corresponding to the Moran process. The state *i* represents all configurations with *i* mutants in the network. The chain goes from state *i* to state *i* + 1 with probability λ*_i_* and to state *i* − 1 with probability *μ_i_*. States 0 and *N* are absorbing states.

The above Markov chain is an example of *absorbing* Markov chains. There is a well known method to calculate *absorption probabilities* and *absorption time* for this kind of Markov chains [10]. It should be emphasized that absorption time differs from fixation or extinction time. In fact absorption time is a weighted average of fixation and extinction times. We will be back to this point later. The heart of this letter is to introduce an analytical method to obtain fixation and extinction times for the above mentioned Markov chain.

Consider a general absorbing Markov chain. The transition matrix of this chain can be written in the following general form which is called *canonical form*:

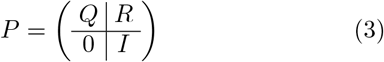

For transition matrix of Eq. (2) we have:

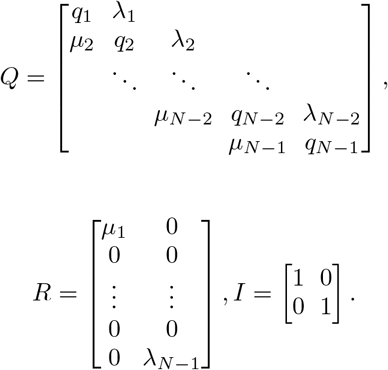

The so-called fundamental matrix corresponding to this transition matrix which is defined as *N* = (*I* − *Q*)^-1^ can be used to calculate absorption probabilities and absorption times. Let us define *t_i_* to be the (average) absorption time of the Markov chain starting from state *i* and 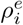 and 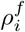 to be extinction and fixation probabilities starting from state *i*, respectively. We emphasize again that by fixation and extinction we mean absorption to the states *S*_0_ and *S_N_* respectively. Following Ref. [10] we use matrix notation to denote these quantities

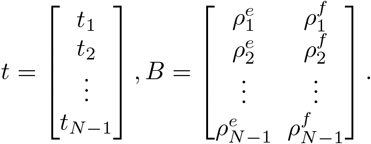

Then, one can easily obtain absorption probabilities and times using the fundamental matrix as *B* = *NR*, *t* = *Nc*, where *c* = (1,1, ⋯, 1)*^t^* is an all-one column matrix.

This method is used to obtain fixation probabilities and absorption time or sojourn time as is called by Hinderson and Traulsen [7]. The point is that here we are interested not only in absorption times but also fixation times and extinction times separately. To this end we need to modify the above general method. In the following we explain the modification that leads to exact formula for fixation and extinction times.

In the transition matrix of the present problem, *R* is a (*N* − 1) × 2 matrix and so is the matrix B. Now, using each column of matrix *B* we define a diagonal matrix and call them extinction and fixation matrices respectively:

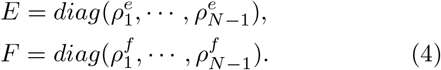

Using the above definitions we define

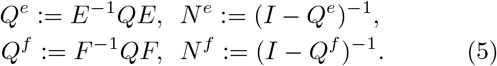

We also denote average extinction and fixation times starting with *i* mutants by 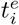 and 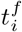, respectively. Then, one can show that these times are obtained as *t^e^* = *N^e^c* and *t^f^* = *N^f^c*, respectively in a matrix form. This can be easily seen for the present Markov chain using the recursive method of Ref. [8], namely by using the recursive relation

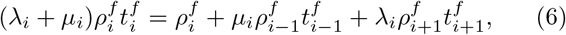

for fixation times and the corresponding one for extinctions times. Our method is applicable for a general absorbing Markov chain with arbitrary number of absorbing states [12].

Applying this method to complete and circle graphs shows complete agreement with the results obtained from recursive equation methods [8, 11]. Generally for graphs whose transition matrices satisfy the condition λ*_i_* = *rμ_i_*, obtaining inverse of tridiagonal matrix (*I − Q*) is straightforward. In these cases the fixation time t1 can be obtained in a closed form

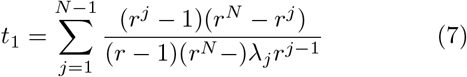

Which is in agreement with the results obtained by other methods [4, 8, 11].

Now we examine this method on a random (Erdös-Rényi) network [13] and calculate the fixation time.

Consider a random network with N nodes where every pair of nodes are connected to each other with probability *p*. For finding fixation time the main step is determining λ_*i*_s and *μ_i_*s. Unlike cycle and complete graphs, here we can not obtain these parameters exactly, therefore we use a mean field approach to obtain them approximately.

Consider a random graph with *N* nodes, *i* number of them are mutants. The probability for a mutant to be selected for reproduction is 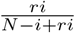. The average nodedegree of a randomly selected node in the Erdཀྵs-Rényi network is approximately *Np*. Now, a natural question is: How many residents are neighbor to a randomly selected mutant? At a first look, one may say there is (*N − i*)*p* residents in the neighborhood of a randomly selected mutant, because there are *N* − *i* residents in the network, each of them may be connected to the selected node with probability *p*. But one should note that the population of mutants grows more of less in a cluster form, therefore it is very likely that the chosen mutant has at least two mutant neighbors. This argument becomes more accurate for smaller values of *p*. This means that the probability that chosen mutant is connected to a resident is 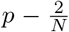 and therefore the average number of residents which is connected to the selected mutant is 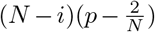. This determines the value of λ*_i_* as

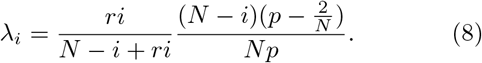

In the same way, one can obtain μ¿

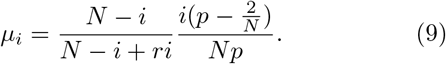

It is obvious from the above relations that in a random graph with this mean-field approximation, the condition λ*_i_* = *rμ_i_* is satisfied and therefore Eq. (7) can be used to calculate the fixation time. Fig. 2 shows the fixation time versus network size for a random network with three different values of fitness. Solid lines and points show analytical an simulation results, respectively. Fig. 3 shows the fixation time versus p in a random graph with size *N* = 100. This figure shows that the fixation time decreases with increasing the connection probability, *p*, and approaches to the fixation time of a complete graph for large values of *p*. This is in agreement with the results of Ref. [9] where exact results for complete graph is reported.

**FIG. 2.**
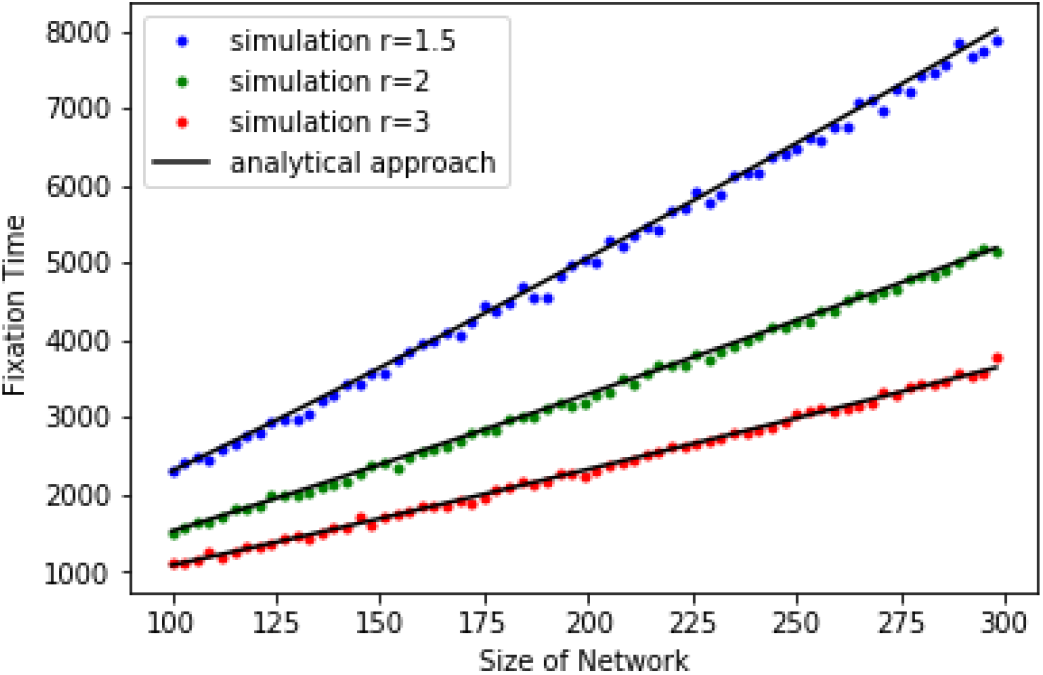
(color on line) Fixation time versus network size of Erdös-Rényi Graph for three different values of fitness. The connection probability is set to *p* = 0.16.

**FIG. 3.**
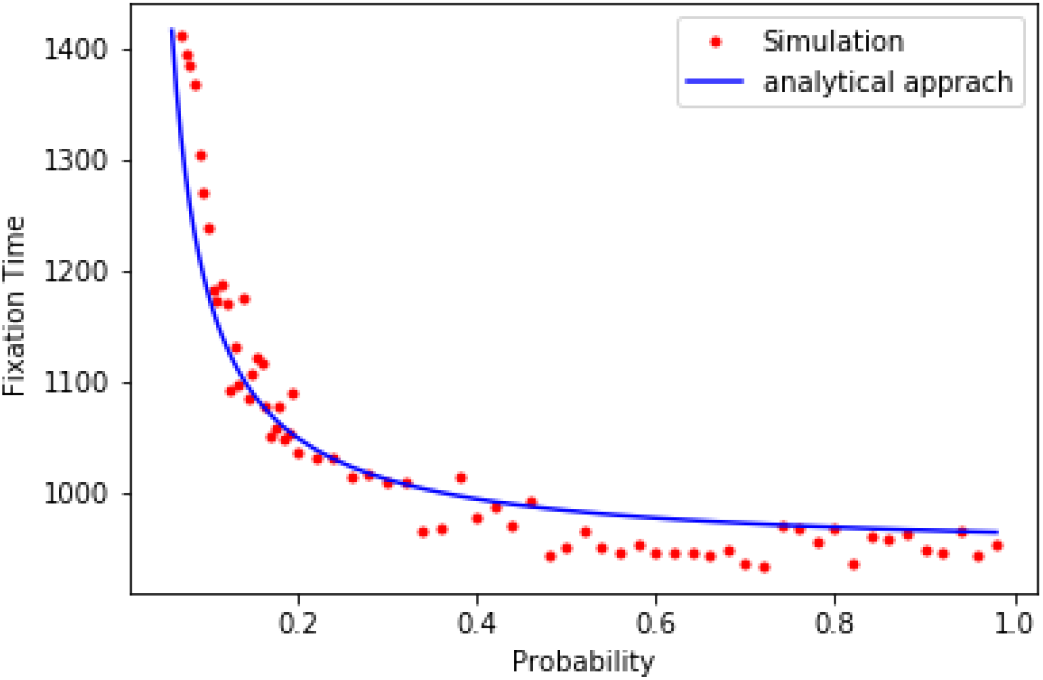
(color on line) Fixation time versus connection probability, *p*, in an Erdös-Rényi Graph with *N* = 100 nodes and fitness *r* = 3.

To summarize we introduced an analytical method to calculate fixation and extinction times of a Moran process on a general evolutionary graph. This method is based on Markov chains and in principle is applicable for all kinds of complex structures. By a modification in the fundamental matrix method we obtained fixation and extinction times separately for an evolutionary process. Results of this method are in agreement with results obtained by recursive equation methods and are confirmed with simulation results for the Moran process on random graphs. This method can easily be used for a dynamical process with more than two absorption states (for example a population with more that two types of species) and provides a straightforward tool to calculate all absorption times.

The authors would like to thank Marzieh Askari for fruitful discussion and comments.

## References

[1] M. A. Nowak, Evolutionary Dynamics, (Cambridge, MA: Harvard University Press 2006).

[2] E. Lieberman, C. Hauert, and M. A. Nowak, Nature (London) 433, 312 (2005).

[3] P. A. P. Moran, Math. Proc. Camb. Philos. Soc. 54, 60 (1958).

[4] P. Shakarian, P. Roos, and A. Johnson, Biosystems 107, 66 (2012).

[5] M. Broom and J. Rychtar, Proc. R. Soc. A 464, 2609 (2008).

[6] M. Frean, P. B. Rainey, and A. Traulsen, Proc. R. Soc. B 280, 20130211(2013).

[7] L. Hindersin and A. Traulsen, J. R. Soc. Interface 11, 20140606 (2014).

[8] M. Askari and K. Aghababaei Samani, Phys. Rev. E 92, 042707 (2015).

[9] M. Askari, Z. Moradi Miraghaei, and K. Aghababaei Samani, J. Stat. Mech. Theory Exp. 073501 (2017). Phys. Rev. E 92, 042707 (2015).

[10] C. M. Grinstead and J. L. Snell. Introduction to Probability, (Providence, RI: American Mathematical Society, 1997).

[11] T. Antal and I. Scheuring, Bull. of Math. Biol. 68,1923 (2006).

[12] K. Aghababaei Samani and M. Hajihashemi, Analytical calculation of absorbing times of absorbing Markov chains, In preparation.

[13] P. Erdös and A. Rényi , Publ. Math. 6, 290 (1959).

